# Human 5’-tailed Mirtrons are Processed by RNaseP

**DOI:** 10.1101/2021.10.15.464553

**Authors:** M. Farid Zia, Jacob Peter, Johnathan Hoover, Kuan-hui E. Chen, Alex Flynt

## Abstract

Approximately a thousand microRNAs (miRNAs) are documented from human cells. A third appear to transit non-canonical pathways that typically bypass processing by Drosha, the dedicated nuclear miRNA producing enzyme. The largest class of non-canonical miRNAs are mirtrons which eschew Drosha to mature through spliceosome activity. While mirtrons are found in several configurations, the vast majority of human mirtron species are 5’-tailed. For these mirtrons, a 3’ splice site defines the 3’ end of their hairpin precursor while a “tail” of variable length separates the 5’ base of the hairpin from the nearest splice site. How this tail is removed is not understood. Here we examine sequence motifs in 5’-tailed mirtrons and interactions with RNA turnover processes to characterize biogenesis processes. Through studying the high confidence 5’-tailed mirtron, hsa-miR-5010, we identify RNaseP as necessary and sufficient for “severing” the 5’ tail of this mirtron. Further, depletion of RNaseP activity globally decreased 5’-tailed mirtron expression implicating this endoribonuclease in biogenesis of the entire class. Moreover, as 5’-tailed mirtron biogenesis appears to be connected to tRNA processing we found a strong correlation between accumulation of tRNA fragments (tRFs) and 5’-tailed mirtron abundance. This suggests that dysregulation of tRNA processing seen in cancers may also impact expression of the ∼400 5’-tailed mirtrons encoded in the human genome.

**SUMMARY:** Abundant non-canonical human miRNAs referred to as tailed mirtrons are processed by RNaseP, which “severs” tail nucleotides to yield a precursor hairpin suitable for Dicer processing. Biogenesis of these miRNAs is correlated with tRFs, which are also products of RNaseP processing.

## INTRODUCTION

microRNAs (miRNAs) are small non-coding ∼22 nucleotide (nt) RNAs processed from stem-loop structures [1]. Biogenesis is initiated by the microprocessor which contains Drosha, an RNase III enzyme, that crops ∼70 nt precursor hairpin from primary transcripts and leaves 2-nucleotide 3’ overhang. The overhangs are recognized by Exportin-5 (XPO5), which carries pre-miRNAs to the cytoplasm for final processing by Dicer, another RNase III, after which they are loaded into effector Argonaute (Ago) proteins. Once recruited by Ago, miRNAs typically base pair with mRNAs via their 5’ nucleotides between position 2-9, inducing degradation or translational repression[2].

While most miRNAs mature through the standard Drosha-Dicer-Ago pathway, numerous additional routes have been identified that take advantage of alternate RNA processing enzymes [3]. Indeed, miRNAs have been found to reside in most major non-coding RNA (ncRNA) varieties such as tRNAs, rRNAs, and snoRNAs. Each of these ncRNAs are cut from longer transcripts, cuts which are repurposed for creation of non-canonical miRNAs. Among these the most prevalent variety is tRFs, which are found in numerous cell types as a sign of stress and are diagnostic of an oncogenic phenotype [4, 5]. This makes many tRNAs dual functional molecules with both tRNA and miRNA capabilities such as tRNA-Ile which can form a 110 bp hairpin encoding hsa-miR-1983 which is also a mature tRNA molecule [6].

Unsurprisingly, the most prevalent RNA processing event in eukaryotes, splicing, also generates miRNAs outside the canonical pathway [7-9]. miRNAs where the hairpin base is defined by splicesome-mediated cuts are called mirtrons and were the first recognized class of non-canonical miRNA. The initial set of mirtrons were minute introns where the precursor fold comprised the length of the entire intron. After splicing, lariat intermediates are linearized by Lariat Debranchase (Ldbr) permitting folding into structures that are suitable for export, Dicer cleavage, and Ago loading. After this discovery, investigation of microprocessor deficient ES cells showed additional types of mirtrons existed where only one side of the hairpin corresponded to a splice site [10]. On the hairpin terminus that was not a splice site a “tail” of nucleotides extended to the scissile base of a distant splice site. Studies in *Drosophila* revealed that removal of a 3’ tail is carried out by the 5’-3’ RNA exosome, which was previously found to be confounded by terminal hairpins on substrates [11]. Critically, this established a paradigm that generic turnover enzymes can participate in productive miRNA biogenesis.

In addition to the alternate method of biogenesis, mirtrons also show a different conservation profile relative to miRNAs. Approximately 80 animal miRNAs are conserved in all metazoans while mirtrons, as well as other non-canonical miRNAs, are rarely found shared at the order taxonomic level [12, 13]. Moreover, mirtron biogenesis modes are not consistently represented. Three types of mirtrons are observed: the original strict mirtrons where splicing generates both ends of the hairpins, 3’-tailed mirtrons where a 5’ splice site coincides with the hairpin base and a tail of nucleotides extends to a distal 3’ splice site, and the 5’-tailed mirtrons where the hairpin base is a 3’ splice site with a tail between the hairpin and a 5’ splice site. Strict mirtrons are found throughout animal genomes with only a handful of examples. Likewise, 3’-tailed mirtrons are rare genomic features, which nevertheless are found in invertebrate and vertebrates at approximately the same rate. Unlike the other mirtron configurations, 5’-tailed mirtrons are extremely abundant in mammalian genomes, while essentially absent or present at the same rate as 3’-tailed mirtrons in lower organisms. Approximately 450 human loci have been identified as sources of diced small RNA duplexes derived from splicing, remarkably 86% are 5’-tailed mirtrons [12, 13].

Other features further separate mirtrons from standard miRNAs. Comparing expression of human canonical miRNAs to mirtrons shows mirtrons have on average several fold lower abundances [12-14]. This is potentially linked to a higher rate of terminal modifications identified on pre- and mature miRNAs that occur on mirtrons [15-17]. Mirtrons are highly modified by untemplated addition of adenine and uridine carried out by noncanonical Poly (A) Polymerases (PAP) or Terminal Uridyltransferase (TUTases) [18, 19]. Nucleotide addition seems to lead to variety of impacts on miRNA biogenesis and functionality; however, effects mostly appear to be inhibitory [18, 20, 21]. Modification can make hairpins poor substrates for Dicer or shift the Dicer cleavage site on the pre-miRNA by altering thermodynamic features and thereby modulating strand selection by Ago [22-24]. Strict and 5’-tailed mirtrons appear to be more prone to uridylation due to the 3’ terminal “AG” of consensus splice sites which is Tailor’s, a terminal uridylyl transferase, preferred substrate. This bias suggests that uridylation of mirtrons is used to specifically suppress their biogenesis. Thus, not only is there selective pressure working to limit mirtron function, but also a targeted modification is applied that further reduces functionality.

Here, we investigate these seemingly contradictory miRNA species through probing biogenesis mechanism of tailed mirtrons in humans. Despite their seeming disfavored biogenesis, mirtrons remain abundant [12, 13]. We show that 3’-tailed mirtrons in mammals likely follows what was found in *Drosophila*. 5’-tailed mirtrons, on the other hand, are produced by the activity of an endoribonuclease, which we identified to be RNaseP. This is the enzyme that is responsible for removal of 5’ leader segments from tRNA precursors, a structure that mirrors the base of 5’-tailed mirtron hairpins [25]. We also find that dysregulation and accumulation of tRFs seen in cancer correlates with a similar accumulation of 5’ tailed mirtrons. This further intertwines processing of 5’-tailed mirtrons with tRNA processing and suggests that the importance of mirtrons might not lie in normal cell physiology, but rather is more impactful during disease processes.

## RESULTS

### 5’-Tailed Mirtrons are characterized by G-quartet containing precursors

To investigate biogenesis of 5’ tailed mirtrons we first sought to identify shared motifs in these miRNAs. To do this, we applied the seqlogo algorithm to the first 20 bases of small RNAs produced from human 5’-tailed mirtron hairpins (Fig 1A-B) [26]. Comparing all species, similar motifs are apparent. 5’ arms are comprised primarily of “G” residues while 3’ arms are typically C and U residues, likely representing intronic polypyrimidine (ppy) tracts. This situation was likewise noted in prior efforts to assess sequence elements in mammalian mirtrons [13]. This suggests that G rich stretches arising adjacent to ppy elements might be sufficient to lead to 5’-tail mirtron formation. Also, the higher confidence G quartet at the beginning of the 5’ arm suggests a major role for hairpin structure in biogenesis of 5’-tailed mirtrons. To confirm this, we assessed the prevalence of G quartets in all human introns relative to 5’ tail mirtron encoding introns, showing significantly higher G quartet occurrence in mirtrons (Fig 1C). Thus, biogenesis of 5’-tailed mirtrons appears linked to RNA structure, specifically polyG tracts. To further connect 5’-tailed mirtron biogenesis to nucleotide content we compared the free energy (ΔG) and GC content of 767 canonical miRNAs and 404 5’-tailed mirtrons (Fig 1D). As a class, 5’-tailed mirtrons show much high GC content relative to most miRNAs. Interestingly, this does not seem to correlate with a substantially different energy profile. If mirtron hairpin energy is greatly divergent from conserved miRNAs this might lead to incompatibilities in Ago loading.

**Figure 1.**
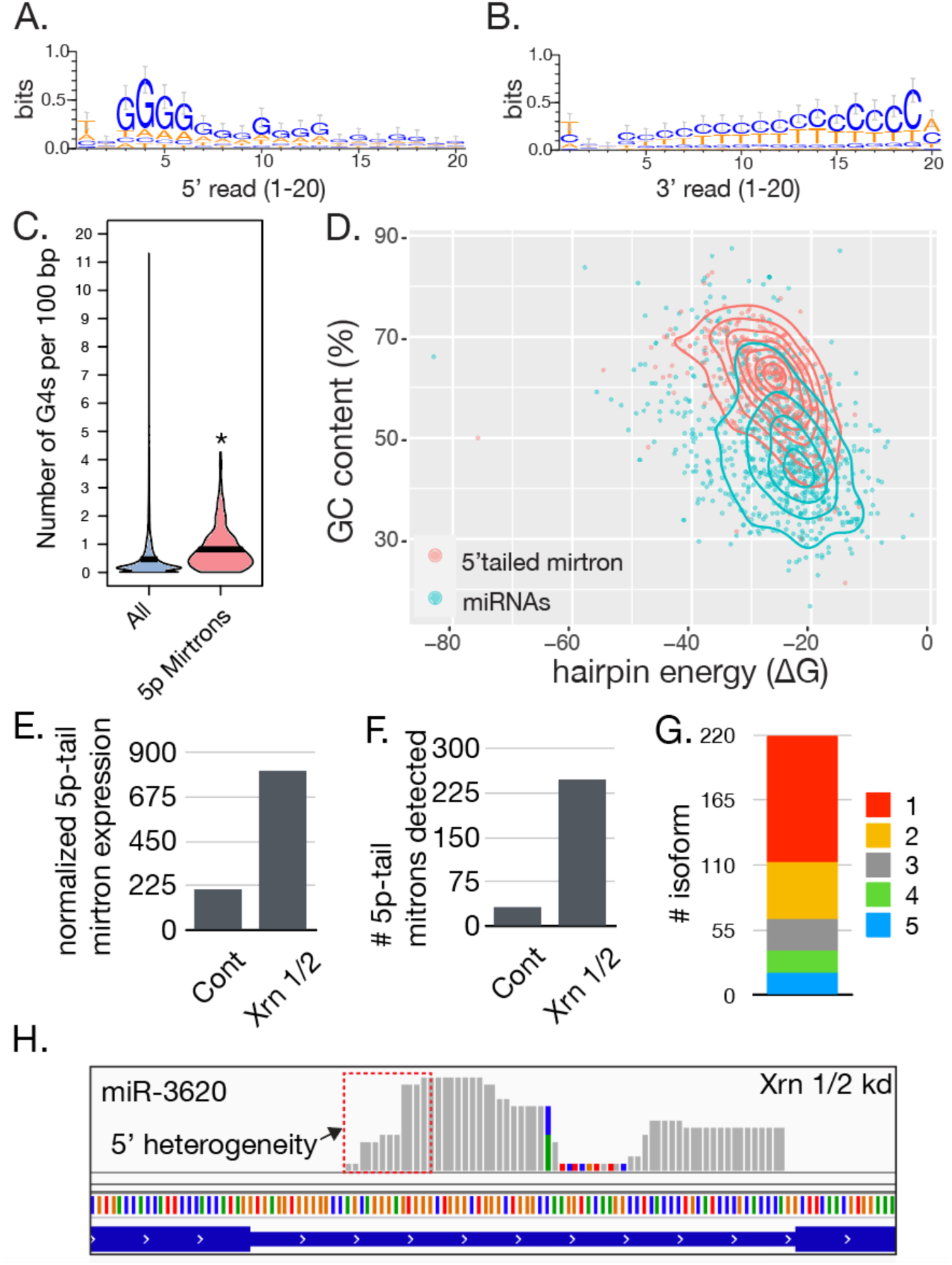
5’-tailed mirtron sequence elements and antagonism by 5’-3’ exoribonuclease. A) Mirtrons Have High “G” content on 5p arms and B) high “CT” content on 3p arms. C) 5’-tailed mirtrons have significantly more G quartets compared to all introns. (*p≤0.05). D) Comparison of ΔG (X axis) and GC (Y axis) contents in 767 canonical miRNAs and 404 5’-tailed mirtrons. Density overlays show separation of the two groups based on GC content, but not ΔG. E) Normalized accumulation of all combined mirtron mapping reads after XRN 1/2 knockdown and in control datasets. F) Number of individual 5’-tailed mirtrons detected in control and XRN 1/2 knockdown libraries. G) Number of isoforms counted for each of the mirtrons only found in XRN 1/2 knockdown libraries. B) Alignment of reads at the miR-3260 locus found in XRN 1/2 knockdown libraries. Red box highlights 5p arm reads differing at their 5’ position.

Considering the bias of 5’ tailed mirtrons for G quartets and high GC content this suggested that part of their biogenesis might be linked to being challenging substrates for RNases. To examine this possibility, we investigated the impact of major 5’-3’ turnover RNases, Xrn1 and Xrn2, in 5’-tailed mirtron production [27]. Using public data from XRN 1/2 depleted cells, 5’-tailed mirtrons are expressed at a significantly higher level compared to control (Fig 1 E-F). A higher level of mirtron expression was observed as well as species that were completely absent in control conditions appeared after XRN1 1/2 knockdown. Initial reporting of these datasets revealed a similar situation, however, this study benefits from a formalized annotation of 5’-tailed mirtrons [13].

These data suggest that exoribonuclease activity antagonizes mirtron production, which led us to investigate the characteristics of mirtrons that only accumulate in Xrn1/2 knockdown datasets. A notable feature of XRN-eliminated mirtrons is the presence of heterogeneous isoforms, where over half were represented by two or more isoforms (fig 1G). This Imprecise processing is consistent with a biogenesis mechanism that is not dedicated to mirtron production, unlike the reliability of Drosha to generate precise hairpin ends. Focusing on a well-expressed mirtron, hsa-miR-3620, greater heterogeneity is seen for 5p arm small RNAs (Fig 1H). Moreover, the greatest isoform diversity is seen at the 5’ end of the 5p read, which is problematic as this shifts the identity of a miRNA’s targets. It also hints that 5’-tail removal is not occurring in a deliberate fashion, suggests a role for XRN activity in limiting 5’-tailed mirtron accumulation, and that this class of miRNA requires features that allow evasion of 5’-3’ exoribounuclease activity.

### Processing of the 5’-tailed mirtron, hsa-miR-5010

To dissect the biogenesis of 5’ tailed mirtrons we examined hsa-miR-5010 which is located in ATP6V0A1 (Sup Fig 1). hsa-miR-5010 is one of the most highly expressed 5’-tailed mirtrons and exhibits relatively consistent 5p arm processing. As a point of comparison, we also examined the mouse 3’-tailed mirtron mmu-miR-668, which is likewise a high confident tailed mirtron (Sup Fig 1) [12, 28]. While there is a homolog of mmu-miR-668 in humans that likewise resides in a large miRNA cluster it is only a 3’-tailed mirtron in mice [28]. Based on the tendency for 5p-tailed mirtrons to harbor polyG tracts we investigated the effect of adding polyG tracts into the tails of hsa-miR-5010 and mmu-miR-668. After each mirtron was cloned into an expression vector, either a 12G element or a mixed identity 20-nucleotide insert was placed in the tail of each mirtron (Fig 2A-C). Wildtype, polyG, and insert constructs were transfected into HEK 293 cells and expression of the mirtron assessed by small RNA sequencing. Consistent with our observations regarding 5’-tailed mirtrons, addition of the polyG tract led to greater accumulation of hsa-miR-5010 reads while the mixed insert did not (Fig 2A-B). hsa-miR-5010 biogenesis was not perturbed by insert, and in all cases terminal modifications were not dramatically altered in ectopically expressed hsa-miR-5010 reads (Fig 2B, Sup Fig 2). The opposite was seen for mmu-miR-668 (Sup Fig3). Both insertions led to decreased expression.

**Figure 2:**
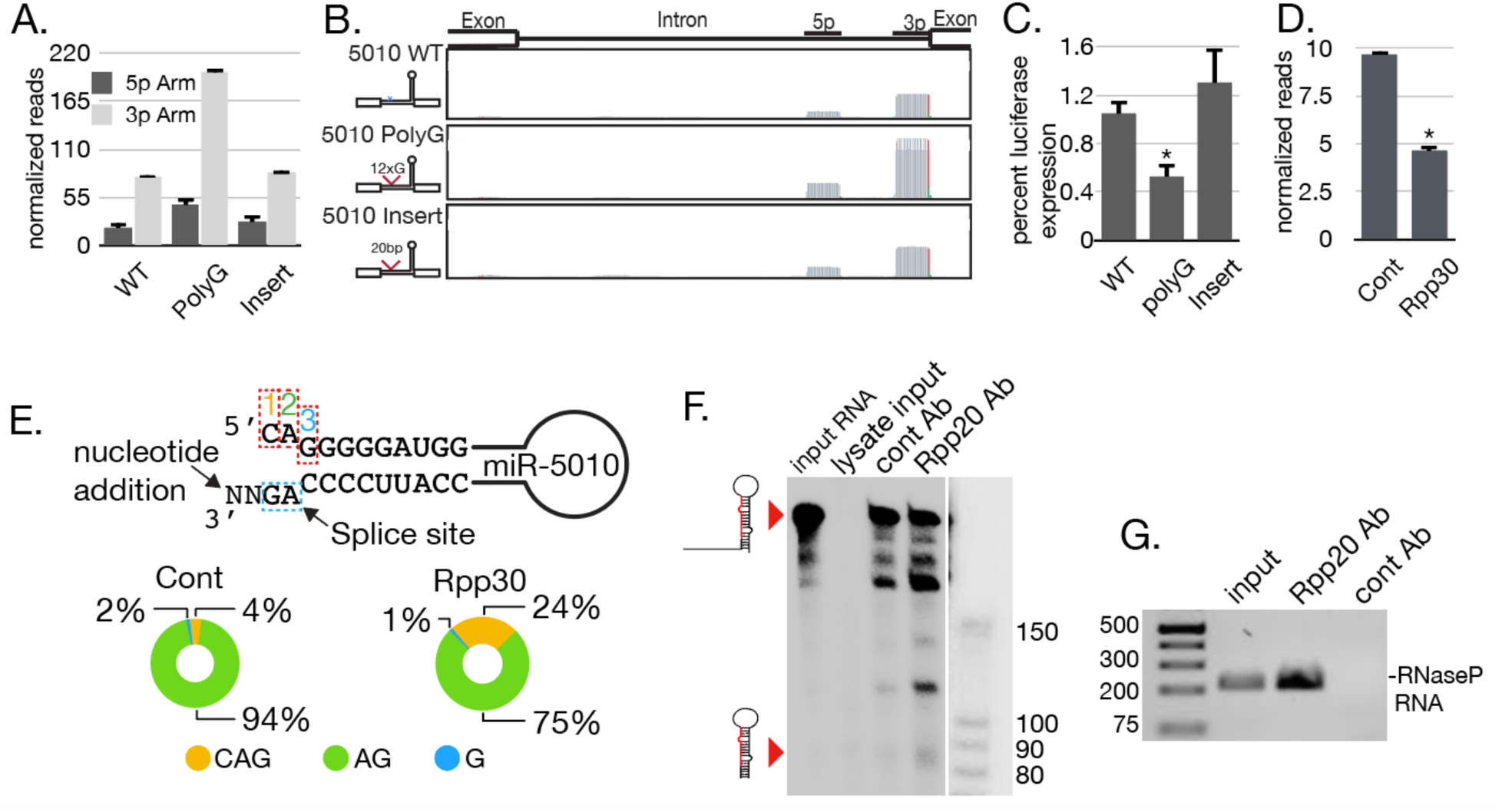
The endoribonuclease RNaseP severs hsa-miR-5010’s 5’ tail. A) Accumulation of expression, determined by small RNA sequencing, from three mirtron constructs (WT, PolyG and Insert) transfected into HEK 293T. Visualization of small RNA read alignments from transfection using the “IGV” software. Left shows the modifications made to the different mirtrons. C) Quantification of luciferase activity from a hsa-miR-5010 sensor following cotransfection with WT, Poly G and Insert hsa-miR-5010 expression plasmids. D) Quantification of hsa-miR-5010 by high through sequencing after transfection of control or Rpp30 siRNAs followed by WT hsa-miR-5010 expression construct. E) Effects of Rpp30 knockdown on 5’ arm heterogeneity. Dominant position found are listed as 1,2,3 as show on the hairpin base diagram. Percent of each found in control and Rpp30 libraries are shown below F) Incubation of radiolabeled hsa-miR-5010 primary intron with immunoprecipitated RNaseP (Rpp20 Ab), mock pull down (cont Ab), or whole cell lysate. Processing products were separated on a PAGE-urea gel. G) RT-PCR of RNaseP complex RNA subunit to verify complex isolation.

Increased hsa-miR-5010 expression after polyG insertion was verified using a luciferase assay to assess target silencing (Fig 2C). For the polyG construct we observed significant reduction in expression, while very little was seen for WT and insert. Together these data suggest that mammalian 5’-tailed and 3’-tailed mirtrons are produced by very different mechanisms. The presence of additional nucleotides in mmu-miR-668 (3’-tailed) resulted in inactivation, which is like what was observed for miR-1017 in drosophila [11]. Inhibition of 3’-5’ exoribonucleases by tail inserts negatively impacted biogenesis. hsa-miR-5010, in contrast, is enhanced specifically by a sequence element inhibitory to exoribonucleases. This is consistent with the effects of Xrn1/2 knockdown on 5’-tailed mirtron expression and suggests that this type of mirtron is produced by endoribonuclease-mediated severing of tails from hairpins.

Considering the structure of tailed mirtrons, where a double-stranded RNA is connected to an unpaired region, it is reminiscent of immature tRNAs. Excision of tRNAs from precursors is carried out by two endoribonucleases: the 5’ Leader cut by RNaseP and the 3’ Trailer by RNaseZ [29]. In the case of 5’-tailed mirtrons the clear fit would be RNaseP. This enzymatic activity is carried out by a multi-subunit complex that coordinates the activity of a ribozyme. To assess the role of RNaseP in mirtron expression, siRNAs targeting the Rpp30 subunit of RNaseP were transfected into HEK cells followed by a second transfection with WT hsa-miR-5010 constructs (Sup Fig 3). After this, small RNA sequencing libraries were created to assess differences in expression and processing. The data shows that the expression of hsa-miR-5010 in Rpp30 knockdown is decreased by half compared to control libraries implicating RNaseP as a factor necessary for 5’-tailed mirtron expression (fig 2D). This led us to evaluate differences in RNAs produced from hsa-miR-5010 in control and Rpp30 knockdown libraries. We detected three different types of endings at the 5’ end of a mature hsa-miR-5010 (fig 2E). Most of the reads (94%) in control started with “AG” (position 2) and when a “U” is added to the 3’ end by uridylation, a 2 nt overhang is obtained which resembles a perfect Drosha product. This is while only 4% of the reads start with “CAG”. On the other hand, in Rpp30 knockdown libraries, 5’ processing has been perturbed and a shift in the first base is observed and the rate of the “AG” is reduced to 75%. This suggests that Rpp30 knockdown has affected the 5’ end processing of hsa-miR-5010. A similar analysis was applied to isoforms differing at the 3’ end but there was no significant difference detected, further implicating Rpp30 in shaping the 5p arm of 5’-tailed mirtrons (Sup Fig 5).

To verify the role of RNaseP in processing hsa-miR-5010, we sought to reconstitute 5’-tail removal using immunopurified complexes (Fig 2F). Antibodies against Rpp20, another RNaseP subunit, were used to isolate complexes. *In vitro* synthesized hsa-miR-5010 primary intron transcripts were incubated with these isolated complexes or with beads bound to mock anti-rabbit IgG antibodies. Incubation with isolates led to significant accumulation of the ∼85 nt hairpin of hsa-miR-5010 in the RNaseP IP, but not the IgG control. In whole lysate incubation, the RNA is degraded by cellular RNases. Immunoprecipitation efficiency was verified by RT-PCR of the RNA subunit of RNaseP, which was greatly enriched in the Rpp20 IP condition relative to control IgG (Fig 2E). These results indicate that not only is RNaseP necessary for processing of hsa-miR-5010, but that it is also sufficient to recapitulate removal of the tail of this 5’-tailed mirtron.

### 5’-Mirtron expression is linked to RNaseP activity

To assess the effects of Rpp30 loss and its correlation with 5’-tailed mirtrons, we compared the expression of all 5’-tailed mirtrons after Rpp30 knockdown to levels found in control libraries (Fig 3A). We find that after knockdown of Rpp30 statistically significant changes in 5’-tailed mirtron expression are almost entirely seen in down regulated species. This is apart from ΔG values which were not predictive of significant down regulation of 5’-tailed mirtrons in Rpp30 knockdown libraries. This reinforces our observation that strength of RNA fold does not typically track with expression. The exception is the handful of outliers, which do exhibit lower ΔG values. However, outliers do not seem to respond consistently to Rpp30 knockdown, suggesting that these changes might be more related to host gene expression changes than instructive of changes in biogenesis.

**Figure 3:**
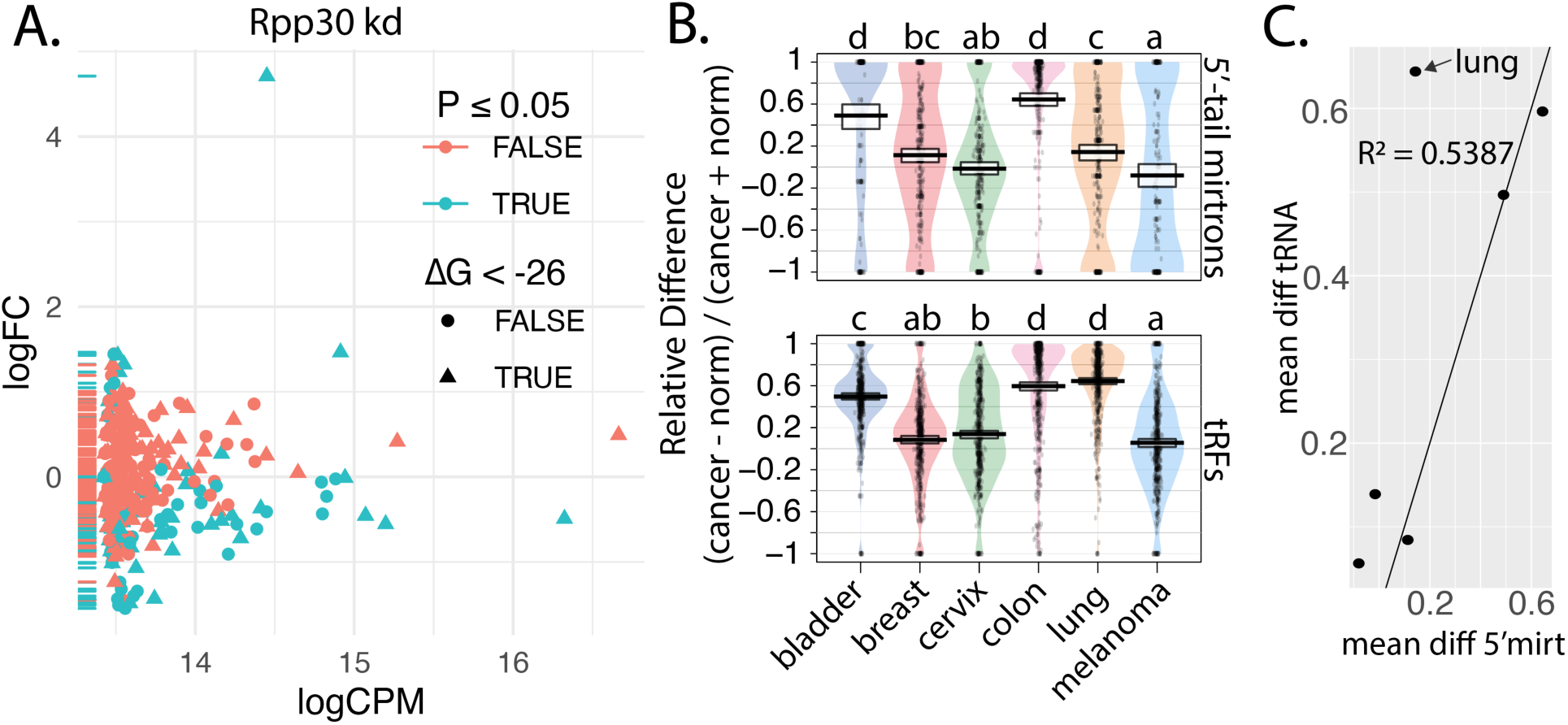
RNaseP processing and global 5’-Tailed Mirtron expression. A) Scatter plot showing expression of 5’-tailed mirtrons following Rpp30 knockdown. Blue colored data points represent statistically significant changes. Red represents non-signficant changes. Data point shape denotes stable (ΔG < -26) or less stable (ΔG > -26). B) Expression of 5’-tailed mirtrons and tRFs in cancers listed on x-axis. Values are relative difference of RPM values (cancer-norm)/(cancer + norm). Letters above plots are significance groups determined by TukeyHSD. D) Plot of Mean relative difference of cancer vs norm in 5’-tailed mirtrons and tRFs plotted by tissues in “B”. R^2^ value indicated.

Given the apparent connection of 5’-tailed mirtrons to RNaseP activity we sought to correlate changes in mirtron expression with biogenesis of tRFs (Fig 3B,C). In small RNA populations, a significant group appear to be derived from pieces of tRNAs [30]. tRFs are not simply degradation products, but are actively loaded into Ago proteins [31-33] Production of tRFs occurs through heterogeneous pathway with most segments of tRNA clover leaf structures giving rise to small regulatory small RNAs [34]. While the function of tRFs is controversial, their expression has been correlated with an oncogenic phenotype [34]. This suggests that cancer cells have a dysregulated tRNA biogenesis, which might also impact 5’-tailed mirtrons. Indeed, after surveying public small RNA sequencing from several cancer types we generally observe greater expression of 5’-tailed mirtrons relative to control tissues (Fig 3B). In all cases the mirtrons were either more abundant in cancer or equally distributed between cancer and norm. Decrease in cancer was not observed. Intriguingly, the degree to which 5’-tailed mirtrons are elevated, tracks almost exactly with the abundances of tRFs. The exception in this situation is lung cancer where mirtrons were enriched to the same degree as tRFs. Even with this outlier, correlation between 5’-tailed mirtrons and tRFs was clear (Fig 3C). These results suggest a compelling correlation between changes in tRNA processing and 5’-tailed mirtrons, reinforcing a shared biogenesis mechanism.

## DISCUSSION

In this study, we identified how a large class of human non-canonical miRNAs, 5’-tailed mirtrons, are processed (Fig 4) [14, 35]. This class of miRNA exhibits distinct features such as high G content on 5’ arms that appears to be a complement to 3’ splice site adjacent ppy tracts. Prior reports on mirtron features report high GC content [15, 36]. Their accumulation appears to be highly antagonized by the activity of 5’-3’ exoribonucleases such that when this enzymatic activity is genetically depleted expression of 5’-tailed mirtrons increases ∼4 fold. Moreover, following XRN 1/2 knockdown 5 times as many mirtron species become detectable. Linked to inhibition of 5’-3’ processing we also observe extreme 5’ arm read heterogeneity. Together these data suggest an antagonistic role for 5’ turnover pathways in 5’-tailed mirtron biogenesis. Consistent with this, insertion of a 12G tract into a high-confidence human 5’-tailed mirtron, hsa-miR-5010, led to a nearly 4-fold increase in mature RNA expression. This is the opposite result to what was observed when testing sequence requirements for miR-1017, a 3’-tailed mirtron encoded in the *Drosophila* genome. For miR-1017, exoribonuclease activity of the RNA exosome was essential for 3’ tail removal. Interestingly, when we perform a similar test with a mammalian 3’-tailed mirtron, mmu-miR-668, a similar result was found. Thus, it would seem both vertebrate and invertebrate 3’-tailed mirtrons likely share a biogenesis mechanism–exoribonuclease processing. They also suggest for mammalian 5’-tailed mirtrons tail removal is likely carried out by an endoribonuclease. Together these results indicate a fundamentally different biogenesis mechanism underlies 5’-tail and 3’-tail mirtrons.

In our studies we implicate the activity of RNaseP in the processing of 5’tailed mirtrons. This ribozyme containing complex has a role in the maturation of tRNAs by removing 5’ leader sequences from precursors. However, studies have found RNaseP to interact and cleave a variety of additional transcripts such as pre-rRNA, snoRNAs, along with additional nuclear RNAs such as the HRA1 antisense RNA [37-39]. Thus, it is not unreasonable to expect this complex to moonlight with mitron primary intron transcripts. Moreover, binding affinity of RNaseP is biases to nucleotide homopolymers, with greatest preference for polyG, a defining characteristic of 5’-tailed mirtrons. This with the tRNA-like single-stranded to double-stranded structure of 5’-tailed mirtrons further reinforces confidence of RNaseP as the likely 5’-tail “severing” enzyme.

One unclear aspect of 5’-tailed biology is their hyper abundance in mammalian genomes. RNaseP is a universal enzyme that is found in all cells and would be able to participate in 5’-tailed mirtron biogenesis. Likewise, as 5’-tailed mirtrons appear to arise from accumulation of polyG tracts adjacent to the universal eukaryotic ppy tract splicing element they should arise at the same rate across many kingdoms [40]. What then leads to the radical accumulation in mammalian genomes? The answer is likely related to the extreme intron length seen in mammals, and the more elaborate alternative splicing patterns [41-43]. We propose two possible scenarios. First, as do the massive length of mammalian introns, exact placement of strong splicing elements such as the ppy are highly favored. A polyG tract could then form through neutral evolutionary process to the point that a viable pre-mirtron hairpin arises, which then become subject to negative selection. Unfortunately, there is significant constraint on the ppy tract such that 5’-tail mitron 3p arms are unable to acquire hairpin disrupting mutations. Harboring the 5’-tailed mirtron is preferable to degrading the strength of the ppy track. In the second scenario, the 5’ arm polyG sequence has a role in modulating splicing as a cis-element. Here by forming a hairpin that occludes the ppy tract, it might lead to alternative 3’ splice choice, and thereby contributing to the greater splicing complexity favored by mammalian gene expression programs. The G-quartets could also be cis-elements that influence alternative splicing through recruitment of hnRNPF [44]. In this second scenario 5’-tailed mirtron expression is an unintended outcome of a different, favorable arrangement.

Considering this case, perhaps 5’-tailed mirtrons can be written off as little more than genomic detritus. They may shape genomes through disfavoring polyG tracts in splice adjacent ppy elements, and thereby apply constraints on ppy neighborhoods. However, it might not be a role in normal cell physiology that should draw the attention of 5’-tailed mirtron function. As potential regulatory molecules that are diverted from Ago loaded by multiple mechanisms such as XRN activity and inhibitory nucleotide addition, these mechanisms could become corrupted to serve the subversive genetics of cancer cells. Indeed, we find a trend towards increased 5’-tailed mirtron expression in cancer-derived small RNA datasets. A similar, and possibly mechanistically related, situation is seen with tRFs, which are also amplified in cancer cells. Further, components of RNaseP are upregulated in nearly all cancer types [45, 46]. Both types of non-canonical miRNAs are RNaseP substrates with unclear roles in regulation of gene expression. They are excellent candidates for exploitation by cancer biology to induce survival promoting gene expression changes as well as excreted RNAs could be used to modulate the function of stromal cells within the tumor microenvironment.

## MATERIALS AND METHODS

### Cell Culture, cloning and transfection

Human embryonic kidney (HEK-293T, ATCC) cells were grown in Dulbecco’s modified Eagle’s medium (DEMEM, GIBCO) supplemented with 10% Fetal Bovine Serum (FBS) and 1% Penicilline-Streptomycine (GIBCO). Cell cultures were maintained at 37° C in 5% CO_2_, 95 % air humidified atmosphere. The constructs to over express the mirtrons: mmu-miR-668 and hsa-miR-5010; were cloned in the pcDNA 3.1 vector. All the clones were transfected into the HEK-293T cells at 90% confluency using FuGene 6 as transfection reagent according to the manufacturer’s instructions (Promega). dsiRNAs targeting Rpp30 (10 nM) were transfected into HEK 293 cells using Lipofectamine 3000 reagent according to the manufacturer’s instructions (Invitrogen). Target are available on www.idtdna.com. Two days after transfection, the total RNA was extracted and used for qRT-PCR or RNA sequencing (Sup Table 1).

### Luciferase Assay

To evaluate functionality of mirtrons constructs a dual reporter luciferase assay was used. Briefly, the hsa-miR-5010 hairpin was cloned downstream of Renilla into the psiCHECK2 vector (Sup Table 1) Then the plasmid constructs (150 ng) were transfected into HEK 293T cells at their 90% confluency using Dual-Luciferase Reporter Assay System (Promega). The Luciferase activity was measured 72 h (48h for DsiRNA and 24 h for plasmid constructs) according to manufacturer’s instructions. Four replicates were used to measure the Luciferase activity and calculated as the ratio of Renilla:Firefly.

### RNA Isolation, qRT-PCR, Sequencing and Analysis pipeline

Total RNA was collected from the transfected HEK cells by Trizol extraction protocol (Invitrogen,Carlsbad, CA) and 2 μg of each sample was used to make small RNA libraries for sequencing. cDNAs was prepared using RevertAid First Strand cDNA Synthesis Kit (Thermoscientific) according to the manufacturer’s recommendation to perform qRT-PCR (BioRad) (Sup Table 1). Sequencing was performed on the Illumina NextSeq platform, and adaptor were clipped using fastx_clipper [47]. Bowtie was used for alignments to hg19 or mm10 [48]. Next, samtools and bedtools were used to quantify reads and examine alignments [49, 50]. The EdgeR R package was used to assess 5’-tailed mirtron expression in libraries [51].

### In Vitro Transcription

Synthetic full length hsa-miR-5010 intron RNA was generated by *in vitro* transcription with MEGAscript® T7 Kit (Invitrogen) using full length intron PCR amplification with a T7 encoding forward primer (Sup Table 1). RNAs were radiolabeled by addition of alpha-P^32^ UTP to the reaction.

### Immunoprecipitation and In Vitro Enzymatic Processing

For immunoprecipitation, the HEK 293T cells were lysed in 0.1% NP-40, 800 nM NaCl, 20 mM Tris at 8.0, 30 mM HEPES, 2 mM Magnesium Acetate, Protease Inhibitor by sonicated. The insoluble fraction was removed by centrifugation at 14,400 rpm for 15 minutes at 4° C. After that the supernatant was incubated for 2 hour with 10 uL of Rpp20 Antibody (Novus Biologicals) immobilized Dynabeads™ Protein G Immunoprecipitation Kit (Invitrogen). The beads were washed five times with IP buffer (0.1% NP-40, 20 mM Tris at 8.0, 30 mM HEPES, 2 mM Magnesium Acetate, 150 mM Potassium Acetate), using a magnetic stand. The enzyme assay reaction was performed at 37° C for 15 minutes with incubation buffer (5% PEG, 10 mM MgCl2, 50 mM Tris-HCl, 100 nM NH4Cl) by incubating RNAs with RNaseP-bound or control IgG-beads. Beads were pelleted and the supernatant was loaded on 8% urea-acrylamide gel and the gel was exposed to phosphor imaging plate (Fujifilm) and was read by Typhoon FLA 7,000. In a separate run, beads were subject to organic extraction followed by RT-PCR of RNaseP RNA subunit.

### Cancer small RNA-metanalysis

Public data from NCBI was used to analyze 5’-tailed mirtron and tRF expression (Sup Table 2). Cancer-norm pairs were from same studies. Alignment and read counting was performed as described above. tRFs coordinates were obtained for hg19 http://gtrnadb.ucsc.edu. Counts were normalized to million of reads mapping (RPM) in the corresponding library. After, relative difference was calculated as (cancer-norm)/(cancer+norm).

---

**Figure 3:**
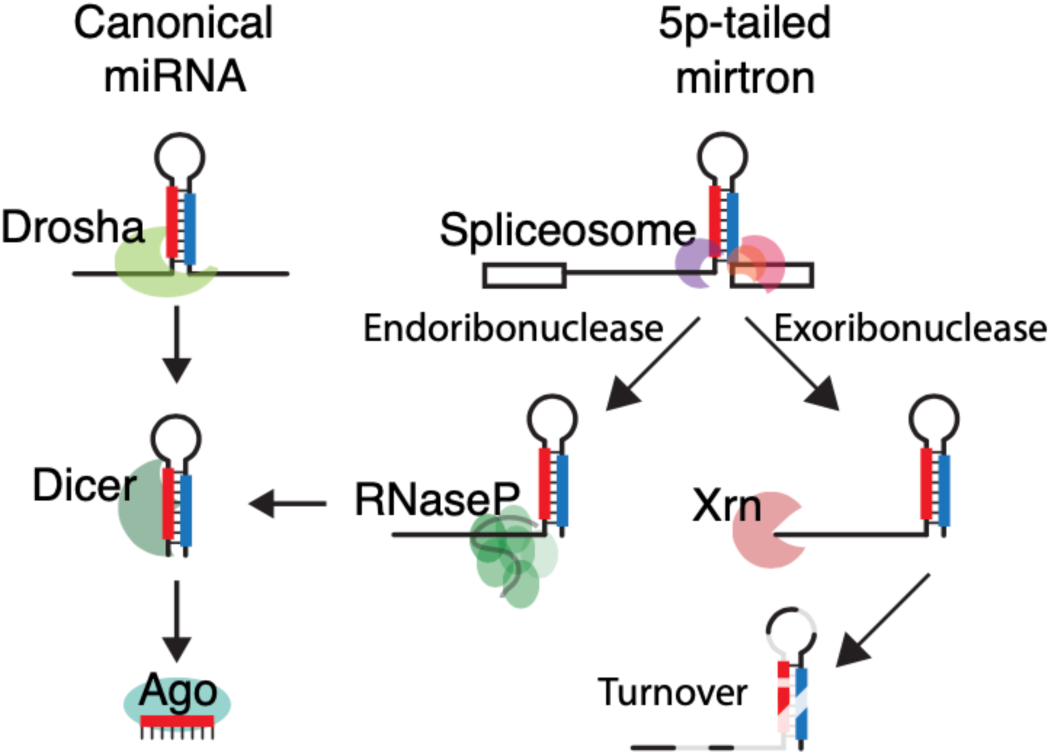
5’-tailed mirtrons can undergo two different processes. They either are processed by RNaseP and produce miRNAs or undergo turnover by XRN exoribonuclease activity and will not be targeted by RNaseP.

## Notes

### Competing Interest Statement

The authors have declared no competing interest.

