## supplement for "Human 5’-tailed Mirtrons are Processed by RNaseP"

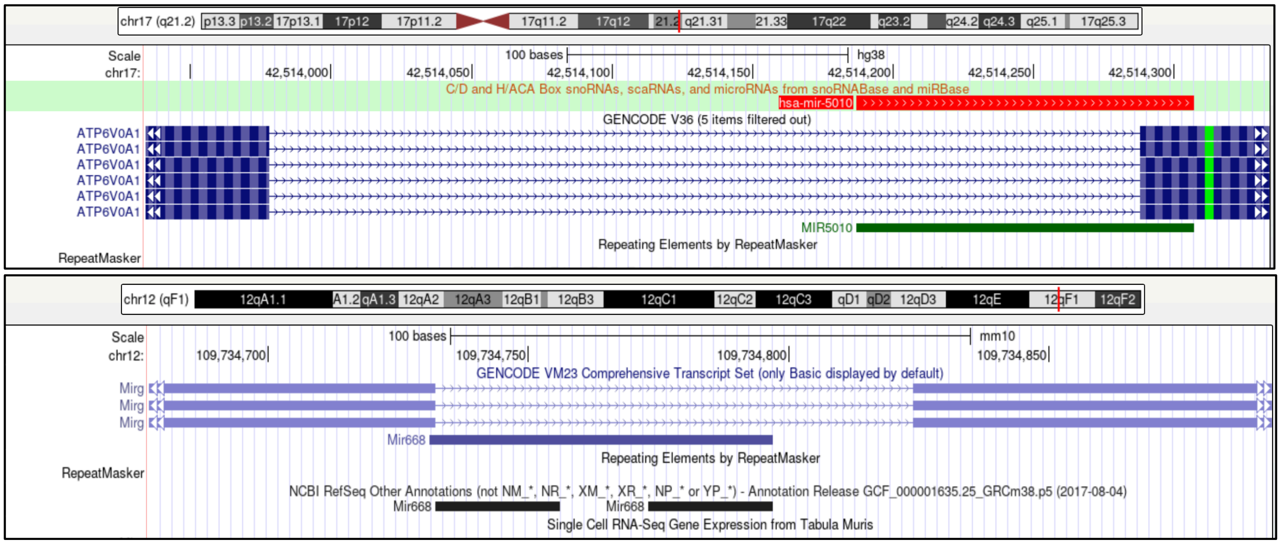


Supplementary Figure 1. UCSC tracks of hsa-hsa-miR-5010 and mmu-mmu-miR-668.

BAR GRAPH SHOWING RATES OF DIFFERENT NUCLEOTIDE ADDITIONS

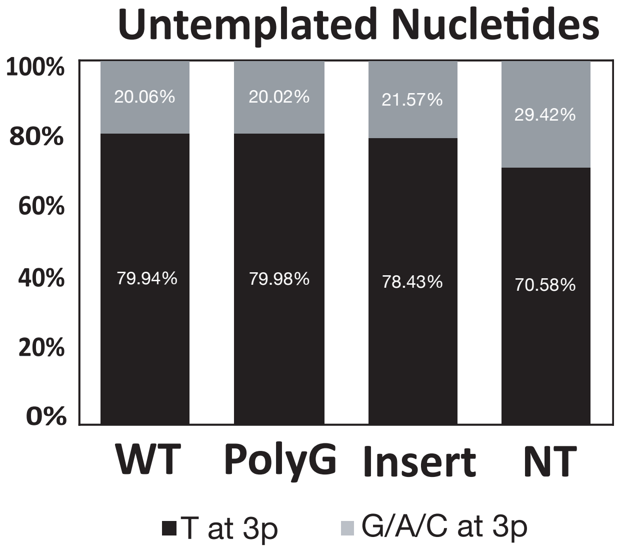


Supplementary Figure 2. Quantification of 3' modifications from transfected hsa-miR-5010 constructs shows similar profiles to endogenous expressed hsa-miR-5010.

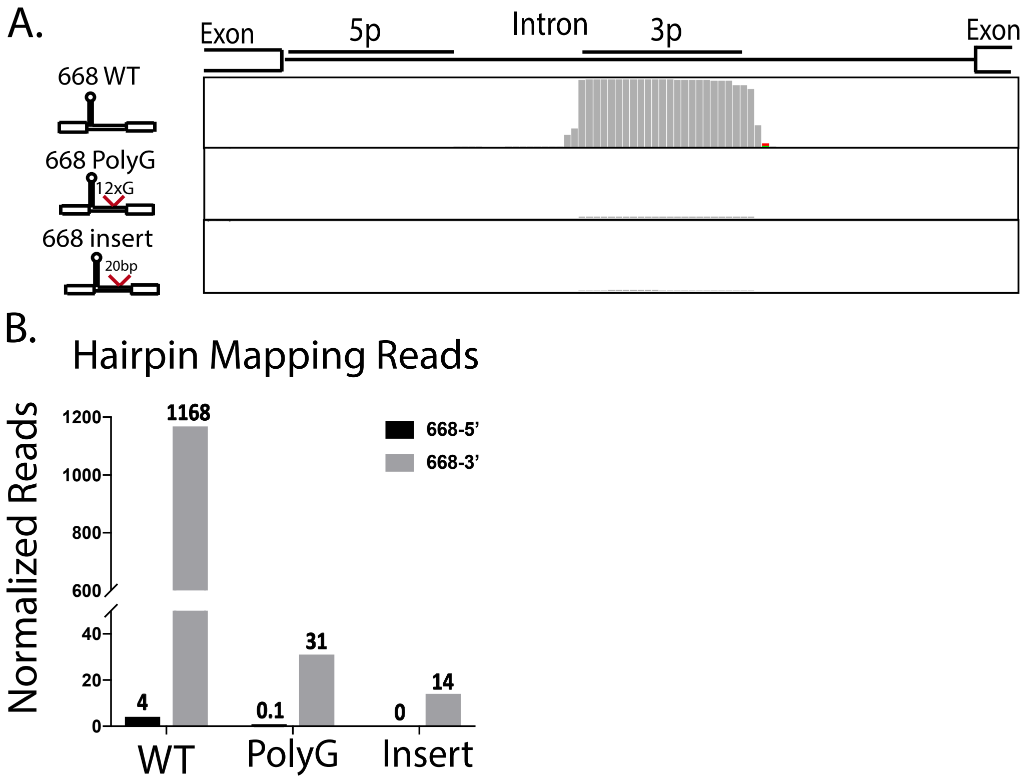


Supplementary Figure 3. A) Three mirtron constructs (WT, Poly G and Insert) were transfected into HEK 293T cells. In the Poly G, 12 "G"s and in the "Insert" 20 random nucleotides were inserted on the 3p tail of mmu-miR-668. Reads from RNA sequencing analysis were visualized using "IGV" software. Mirtrons with Poly G tract showed higher mature mirtrons. Untemplated "T" nucleotide reads at the 3' end are observed mostly on 3p end of the 3p arms but less than that in 5'-tailed mirtrons. B) Hairpin mapping reads were normalized based on the total number of reads in each library. WT showed much higher matured mirtrons.

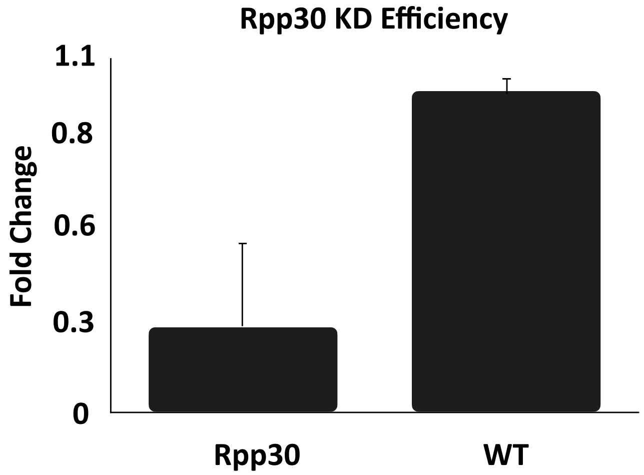


**Supplemental Figure 4**. Knockdown efficiency of Rpp30 dsiRNAs.

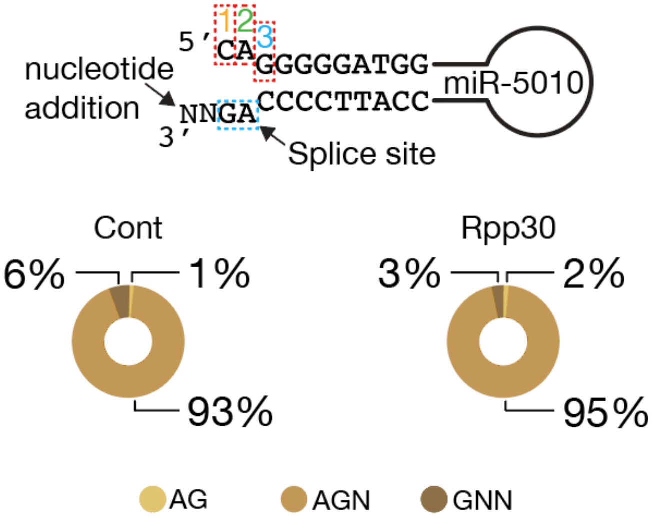


**Supplemental Figure 5.** No change in hairpin 3’ residue identity seen after Rpp30 knockdown. Single nucleotide addition, which when combined with cleavage at the “2” 5’ position yields an ideal Drosha product mimic.

Supplementary Table 1

| Construct | Sequence |
|---|---|
| RPP30 F | tcagcatggcggtgtttgcag |
| RPP30 R | gcggctgtctccacaagtccg |
| B-Actin F | catgtacgttgctatccaggc |
| B-Actin R | ctccttaatgtcacgcacgat |
| hsa-miR-5010 T7 F | taatacgactcactataggTGAGTACCTCTCTCCGGGC |
| hsa-miR-5010 R | ctggggaatgggagacacaaaatcc |
| mmu-miR-668 F | gtaagtgtgcctcgggtgag |
| mmu-miR-688 R | cattcacacggagcccactc |
| hsa-miR-5010 + Strand, Luciferase insert | ggccgcCCATCC CCCactgacCCATCCCCCactgacCCATCCCCC actgacCCATCCCCCc |
| hsa-miR-5010 - Strand, Luciferase insert | tcgagGGGGGATGGgtcagtGGGGGATGGgtcagtGGGGGA TGGgtcagtGGGGAT |
| dsiRNA used | hs.Ri.Rpp30.13.1 |
| dsiRNA used | CD.Ri.195363.13.5 |
| Rpph1F | ccactgatgagcttccctcc |
| Rpph1F | ggaggagagtagtctgaattgg |

Supplementary Table 2

| Cancer | Cancer | norm |
|---|---|---|
| Lung | SRR14634327,SRR14634347,SRR14634359 | SRR14634273,SRR14634279,SRR14634283 |
| Melanoma | SRR14510928,SRR14510929,SRR14510930 | SRR14510925,SRR14510926,SRR14510927 |
| Bladder | SRR333655,SRR333657,SRR333659 | SRR333656,SRR333658,SRR333660 |
| Colorectal | SRR8932122,SRR8932124 | SRR8932123,SRR8932125 |
| cervical | SRR11095745,SRR11095746,SRR11095747, SRR11095748 | SRR11095741,SRR11095742,SRR1109574, SRR11095744 |
| Breast | SRR8330380,SRR8330384,SRR8330374, SRR8330376,SRR8330378,SRR8330382 | SRR8330375,SRR8330377,SRR8330379, SRR8330381,SRR8330383,SRR8330385 |
